# Structural analysis of the interaction between the bacterial cell division proteins FtsQ and FtsB

**DOI:** 10.1101/362335

**Authors:** Danguole Kureisaite-Ciziene, Aravindan Varadajan, Stephen H. McLaughlin, Marjolein Glas, Alejandro Montón Silva, Rosa Luirink, Carolin Mueller, Tanneke den Blaauwen, Tom N. Grossmann, Joen Luirink, Jan Löwe

## Abstract

Most bacteria and archaea use similar proteins within their cell division machinery, which uses the tubulin homologue FtsZ as its central organiser. In Gram-negative *Escherichia coli* bacteria, FtsZ recruits cytosolic, transmembrane, periplasmic and outer membrane proteins, assembling the divisome that facilitates bacterial cell division. One such divisome component, FtsQ, a bitopic membrane protein with a globular domain in the periplasm, has been shown to interact with many other divisome proteins. Despite its otherwise unknown function, it has been shown to be a major divisome interaction hub. Here, we investigated the interactions of FtsQ with FtsB and FtsL, two small bitopic membrane proteins that act immediately downstream of FtsQ. In biochemical assays we show that the periplasmic domains of *E. coli* FtsB and FtsL interact with FtsQ, but not with each other. Our crystal structure of FtsB bound to the β domain of FtsQ shows that only residues 64-87 of FtsB interact with FtsQ. A synthetic peptide comprising those 24 FtsB residues recapitulates the FtsQ:FtsB interactions. Protein deletions and structure-guided mutant analyses validate the structure. Furthermore, the same structure-guided mutants show cell division defects *in vivo* that are consistent with our structure of the FtsQ:FtsB complex that shows their interactions as they occur during cell division. Our work provides intricate details of the interactions within the divisome and also provides a tantalising view of a highly conserved protein interaction in the periplasm of bacteria that is an excellent target for cell division inhibitor searches.

**Importance:** Cells in most bacteria and archaea divide through a cell division process that is characterised through its filamentous organiser, FtsZ protein. FtsZ forms a ring structure at the division site and starts the recruitment of 10-20 downstream proteins that together form an elusive multi-protein complex termed divisome. The divisome is thought to facilitate many of the steps required to make two cells out of one. FtsQ and FtsB are part of the divisome, with FtsQ being a central hub, interacting with most of the other divisome components. Here we show for the first time how FtsQ interacts with its downstream partner FtsB and show that mutations that disturb the interface between the two proteins effectively inhibit cell division.

## Introduction

The divisome is a macromolecular complex formed by at least 12 essential proteins and even more non-essential proteins. It effects bacterial cell division through a number of processes, including cell constriction, synthesis of the septal peptidoglycan (PG) wall, and ultimately cell separation (1–5). In *Escherichia coli*, divisome assembly starts with formation of a ring structure localised at midcell, containing the bacterial tubulin-homologue FtsZ in the cytoplasm (6, 7) and anchoring of the ring in the inner membrane by FtsA and ZipA (8, 9). This is followed by recruitment of further cell division proteins FtsEX, FtsK, FtsQ, FtsL, FtsB, FtsW, FtsI, and FtsN in order of their localisation inter-dependence, all of them trans-membrane proteins. The functions of several of these divisome proteins have been deduced, such as the role of FtsEX in transmembrane regulation of septal PG hydrolytic enzymes (10), FtsK’s role in XerCD-mediated chromosome decatenation (11) and FtsW/PBP1b/FtsI’s proposed role as hybrid septal peptidoglycan synthase with transglycosylase and transpeptidase activities, respectively (12). FtsN has a particularly interesting role as the trigger for septal peptidoglycan synthesis, depending on the assembly of the entire complex, somehow affecting FtsA on the cytoplasmic side of the cell membrane directly as a feedback or even checkpoint mechanism (13). FtsQLB, FtsI and FtsN have only single transmembrane helices anchoring them in the membrane (bitopic membrane proteins) in addition to globular periplasmic domains, where they act on/react to PG synthesis.

FtsQ is considered to play a central, yet enigmatic role in assembly of the divisome through a multitude of interactions as no enzymatic function is known for this protein (14). Two-hybrid analyses have suggested that FtsQ interacts with ~10 cell division proteins of which the interactions with FtsB and FtsL were confirmed biochemically (15). The FtsQBL complex may form independently before its recruitment to midcell by FtsK, where it interacts with the later divisome proteins needed for cell division (16).

FtsQ is a particularly attractive target for the development of inhibitors of protein-protein interactions (PPIs) that block bacterial division because (i) it is an essential protein of low abundance (~50-250 copies per cell) (17) (ii) it has multiple interactions in the much more easily accessible periplasm (14, 16), and (iii) it has no obvious eukaryotic homologues (18). Previously, the structure of the periplasmic domain of FtsQ from *E. coli* has been solved by X-ray crystallography (19) and shown to consist of two subdomains, named α and β. Together with the trans-membrane domain (TMD, a single bitopic helix in FtsQ), the α domain is believed to be required for recruitment by FtsK although other interactions have been ascribed to this domain as well (19, 20). The α domain is located directly downstream from the TMD in the sequence of FtsQ and corresponds to the more broadly distributed POTRA domains that have been implicated in transient PPIs in a range of transporter proteins (21). The β domain has been implicated in multiple interactions including those with FtsB and FtsL (19).

FtsB and FtsL are small bitopic inner membrane proteins (103 and 121 residues in *E. coli*, respectively). FtsB and FtsL have also been suggested to form a distinct subcomplex prior to localisation to the septum making a strictly sequential recruitment less likely. Both proteins may contain a leucine-zipper motif in their periplasmic domain that together with the TMDs have prompted suggestions that FtsB and FtsL form a heterodimeric or tetrameric complex (22, 23). Recently, an *in vivo* scanning photo cross-linking approach to map interactions of FtsQ with FtsB and FtsL at the amino acid level has been conducted, considering one fifth of all surface exposed residues (24). Two hotspots for the interactions with FtsBL were identified on FtsQ: one in the α domain close to the membrane around residue FtsQ R75, primarily with FtsL, and a more pronounced hotspot in the conserved distal part of the β domain around residue FtsQ Y248, primarily interacting with FtsB. In addition, it was previously shown with purified proteins *in vitro* that artificially dimerised FtsB and FtsL, replacing their TMDs with a heterodimeric coiled coil fragment, bind to the periplasmic portion of FtsQ (25). In those experiments, the periplasmic domain of *E. coli* FtsQ was co-purified with the heterodimerised *E. coli* FtsB and FtsL constructs as a stable trimeric complex. FtsB was also shown to interact with FtsQ in the absence of FtsL, whereas FtsL on its own failed to co-purify with FtsQ.

Here we present further quantitative biochemical investigations of the periplasmic interaction between FtsQ, B, L and FtsN, showing that the FtsQB complex is formed with a sub-micromolar dissociation constant as determined by surface plasmon resonance (SPR). SPR does not indicate interactions between FtsB and FtsL but shows a trimeric FtsQLB complex. FtsN interacts independently of FtsB with FtsQ, whereas the interaction of FtsQ with FtsL is not additive with FtsB. In line with these findings, we present a crystal structure of the periplasmic domain of *E. coli* FtsQ in complex with the periplasmic domain of *E. coli* FtsB. The structure resolves residues 64-86 of FtsB that form an α helix and a β-strand, linked by a loop. The region in FtsQ that interacts with FtsB has Y248 at its centre and is at the membrane-distal end of the protein, which is in agreement with the previous crosslink data (24). Mutational analysis coupled with cellular microscopy and also SPR confirmed residues in the interacting region highlighted by the crystal structure that are critical for binding of FtsB to FtsQ, and consequently for functioning of these proteins in cell division.

## Results

### The periplasmic domain of FtsB binds to FtsQ, but not FtsL

To understand the periplasmic interactions of FtsB with FtsQ and FtsL, we performed surface plasmon resonance (SPR) experiments with purified protein domains (Figure 1A & C). The periplasmic domain of *E. coli* FtsB comprising residues 22-103 binds strongly to *E. coli* FtsQ (residues 58-276) immobilised on the SPR chip, with a major K_d_ of 0.8 µM. Equally, the periplasmic domain of FtsL binds to FtsQ, however in this experiment FtsL was immobilised and the binding and release were so fast that parameters could not be determined reliably. Surprisingly, when FtsB was added to immobilised FtsL, no binding could be detected. To investigate this further, FtsB and FtsQ were added together to immobilised FtsL, resulting in measurable binding with a K_d_ of 11 µM. The lack of binding of FtsL alone to FtsB is surprising because it has been proposed that FtsB and FtsL form a coiled coil/leucine zipper complex with each other and when binding to FtsQ (22, 23). When analysing the propensity of FtsL and FtsB to form coiled coils with COILS (Figure 1D) (26), only *E. coli* FtsB showed significant coiled coil content between residues 29-70 (or 77). 2ZIP leucine zipper predictions (27) were also negative for FtsL but positive for FtsB (not shown). We think given these data it is unlikely that FtsB and FtsL form a 1:1 coiled coil/leucine zipper complex and we could detect no binding of their periplasmic domains biochemically. However, FtsQ and FtsB together do bind to FtsL and the binding is not additive in the sense that if FtsL and FtsB bound independently to FtsQ, the resulting binding would be stronger than their individual binding, which is not the case. To test this further, we investigated the binding of the periplasmic domain of FtsN to immobilised FtsQ as a control (Figure 1B & C). *E. coli* FtsN comprising residues 57-319 was bound to immobilised FtsQ using SPR and it bound with a K_d_ of 2.5 µM. When FtsB and FtsN were bound to FtsQ together, a tighter K_d_ of 0.12 µM resulted, and the resulting binding could be simulated by assuming additive, that means independent binding of FtsB and FtsN, as shown by the calculated plot in Figure 1B. We conclude that FtsB and FtsL alone bind well to FtsQ, and as a complex FtsB and FtsL bind to FtsQ dependent on each other, not binding to their own, independent binding sites on the surface of FtsQ and/or inducing a conformational change in FtsQ. In contrast, the binding of FtsN and FtsB to FtsQ is independent of each other hence additive. Surprisingly, the periplasmic, soluble domains of FtsB and FtsL do not bind to each other in isolation and only FtsB is predicted to form a coiled coil, presumably with itself.

**Figure 1.**
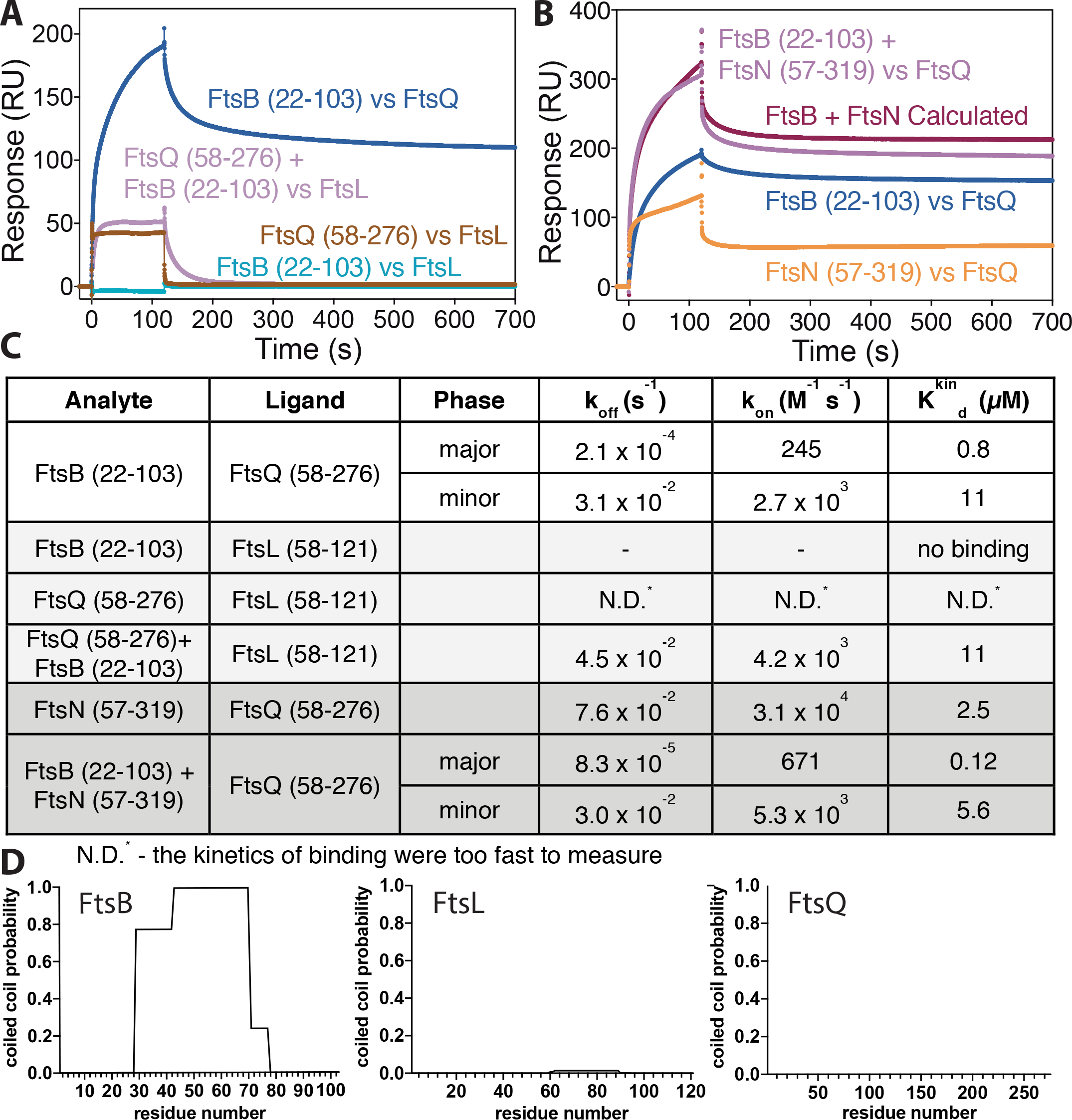
The periplasmic domains of *E. coli* FtsQ and FtsB form a stable complex *in vitro*. A) Surface plasmon resonance (SPR) experiments investigating the interactions of the periplasmic domains of *E. coli* FtsQ, FtsB and FtsL. FtsB binds to FtsQ. FtsL binds to FtsQ, and FtsL and FtsB together also bind to FtsQ, although synergistically, not independently. FtsB and FtsL do not bind to each other in isolation. Note that the proteins mentioned last were immobilised. B) SPR investigation of the interaction of the periplasmic domain of *E. coli* FtsN with FtsQ. A control, showing that binding of FtsB and FtsN to FtsQ is additive, meaning they bind independently to FtsQ, different to how FtsB and FtsL bind. C) Summary table showing quantitative analysis of the SPR data presented in panels A) and B). D) Coiled coil predictions of full-length *E. coli* proteins FtsB, FtsL and FtsQ, calculated with COILS (26). Note that only FtsB shows significant coiled coil prediction, between residues ~29-70 (or 77). This makes it unlikely that FtsB and FtsL form a canonical heteromeric coiled coil and we show in panel A that they do not interact *in vitro* on their own. Please note that also predictions with 2ZIP (27) were negative for FtsL (but positive for FtsB) (not shown).

### Crystal structure of the periplasmic FtsQ:FtsB complex

Inspired by the SPR results, we purified the complex between the periplasmic domains of *E. coli* FtsB (residues 22-103) and FtsQ (residues 58-276). The final size exclusion chromatogram of the purification is shown in Figure 2A, showing two peaks. Both peaks yielded the same crystals and peak A, of unexpected high apparent molecular weight is presumed to be a dynamic oligomeric state of the complex. Both peaks A and B elute as a defined complex with 1:1 stoichiometry (Figure 2B). We obtained tetragonal crystals of the complex and solved the X-ray crystal structure to 2.6 Å resolution by molecular replacement with a previous *E. coli* FtsQ structure (PDB ID 2VH1, Table 1) (19). The resulting structure shows the previously reported two-domain architecture of FtsQ (α and β domains) largely unchanged. FtsB binds to the C-terminal β domain of FtsQ, presumably furthest away from the membrane (Figure 2C & 5B). Only a C-terminal portion of FtsB was resolved in the crystals, comprising residues 64 to 87 (22-103 were present). FtsB forms a helix, followed by a linker largely in β-strand conformation, leading to the final β-strand that extends the central β-sheet of the FtsQ β domain by binding to its last β-strand (Figure 2C). Analysis of the binding mode of FtsB to FtsQ reveals a number of key interactions that are depicted in the stereographic Figure 2D. The only helix resolved in FtsB, approximately residues 65-75, makes a number of charged interactions with the rest of FtsB and FtsQ, most notably two salt bridges that may be involved in stabilising FtsB’s conformation (E68-R79 and R72-E82). Both salt bridges are also involved in binding to FtsQ via D245 and R247. Further charged or polar interactions with FtsQ are mediated by FtsB E65, E69 and N73 (not highlighted in Figure 2D). In the crystals, the resolved part of FtsB forms a tight dimer, the significance of which we have not investigated (Figure 2E). Figure 2E also shows the phased anomalous difference density from a SAD experiment with selenomethionine substituted M77, validating the model building, in order to produce absolute certainty for the chain trace since the resolved FtsB domain is small.

**Figure 2.**
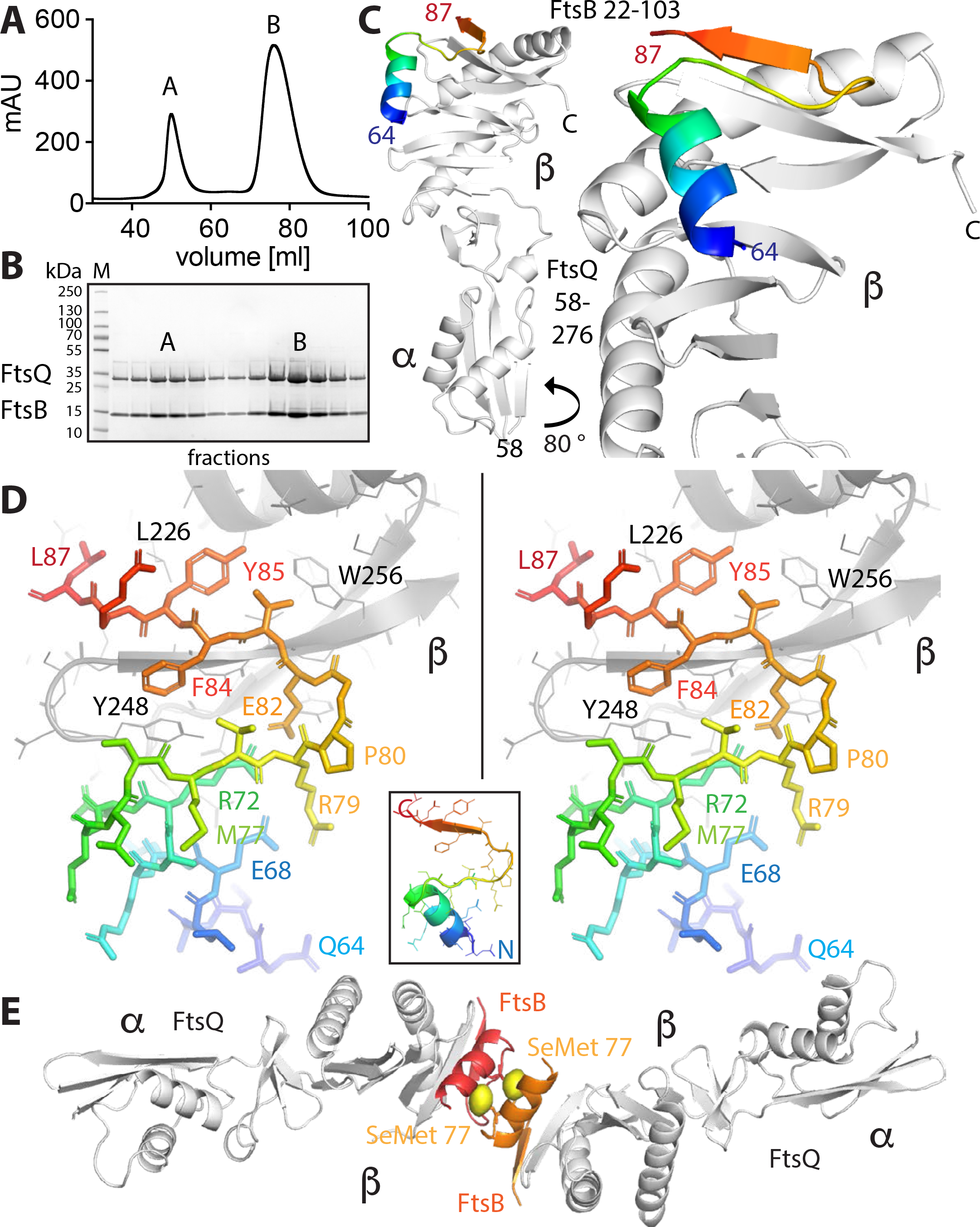
Crystal structure of the complex between the periplasmic domains of *E. coli* FtsB and FtsQ. A) Size exclusion profile showing elution of the FtsQB complex. The complex elutes as two peaks (see Figure 5A and Suppl. Figure S6A for SEC-MALS and AUC data on the same complex) that both produced crystals and are most likely related to oligomerisation or dimerisation of FtsB. C) Crystal structure of the complex determined to 2.6 Å resolution by molecular replacement. Crystallographic data are listed in Table 1. Only residues 64 to 87 of FtsB are resolved in the structure. FtsB forms a short helix, a connecting loop and a β-sheet that aligns in an antiparallel orientation with the last strand of the β domain of FtsQ. D) Stereo plot of FtsB 64-87 in stick representation, showing key residues involved in interactions with FtsQ, and also internal contacts that are important for FtsB to adopt this particular structure. Coloured from N- to C-terminus blue to red. Inset shows the same orientation as a ribbon plot. E) In the crystals, FtsB forms a tight dimer that buries hydrophobic residues, including methionine 77. To be certain about the register of the amino acids of FtsB, we replaced M77 with SeMet and performed a SAD experiment (Table 1). The resulting phased anomalous difference density highlights the only methionine in FtsB in the correct location, validating our interpretation.

**Table 1.**
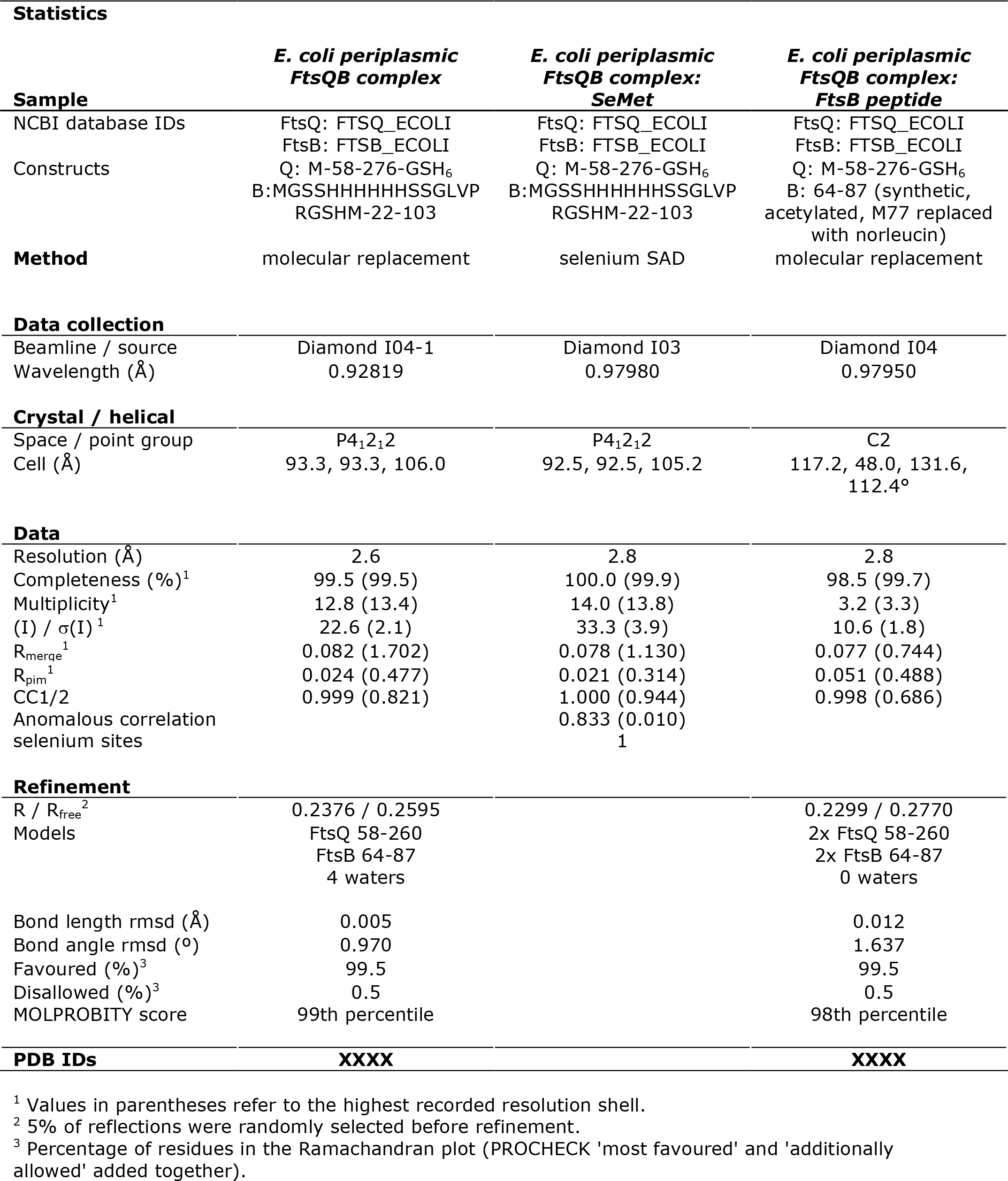
Crystallographic data

### FtsB(64-87) is necessary and sufficient for the FtsQ:FtsB interaction

First, we aimed at an *in vitro* validation of the crystal structure by investigating structure-guided mutant FtsB proteins and their binding to FtsQ (Figure 3A & C). We chose FtsB R72A that forms a salt bridge within FtsB with E82 and is in direct contact with F84A that is positioned next to Y248 on the surface of FtsQ (Figure 2D). Both mutant FtsB proteins showed approximately ten-fold reduced binding to immobilised FtsQ. Even more convincingly, FtsQ Y248W, a somewhat conservative mutation, showed no binding whatsoever when FtsB was tested. FtsQ Y248 (with A253) forms the central hydrophobic patch that FtsB latches onto (Figure 2D). All mutants tested confirmed the binding mode of FtsB to FtsQ as shown by the crystal structure of the complex (Figure 2) and this is further supported by an analysis of sequence conservation across FtsB homologues as depicted in Suppl. Figure S1 that shows strong conservation for residues shown in the crystal structure to interact with FtsQ.

**Figure 3.**
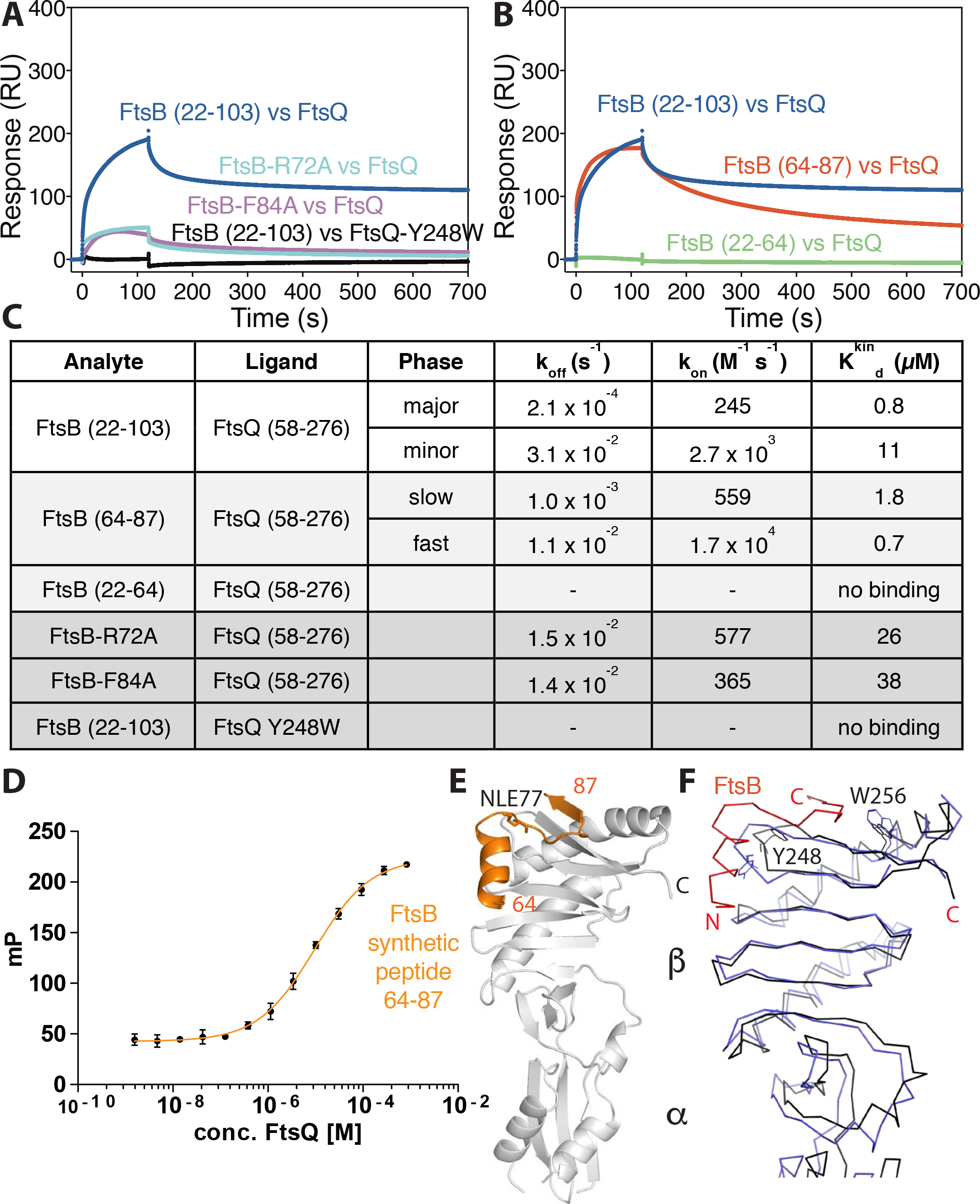
Validation of the crystal structure. A) SPR experiments showing that FtsB mutants R72A and F84A display significantly compromised binding to FtsQ, as does FtsQ mutant Y248W to FtsB, which showed no binding. The crystal structure of the complex of FtsB and FtsQ implicates FtsB F84 and FtsQ Y248 in forming the binding interface. FtsB R72 is involved in a key salt bridge with FtsB E82 and its interruption seems to abrogate binding between FtsB and FtsQ as well. Note that these and other mutants were also investigated *in vivo* as described in Figures 4 and Suppl. Figures S3-S5. B) SPR experiments investigating the role of the FtsB residues that were not resolved in the crystal structure, as only residues 64-87 were visible, 22-63 were not. FtsB binding to FtsQ only requires residues from 64 onwards until 87 and it is to be concluded that the remainder of the protein in the crystals is disordered. C) Summary table quantifying the SPR data in panels A) and B). D) Corroborating the point that only FtsB residues 64-87 are needed for the interaction between FtsB and FtsQ, a fully synthetic peptide was produced (see Suppl. Figure S2) and its binding to FtsQ investigated by fluorescence polarisation as the peptide also carried an FITC moiety at the N-terminus. Fitting of the binding curve yielded a K_d_ of 9.5 µM, similar to the values obtained with recombinant FtsB 22-103 and 64-88 in B) and C). E) In fact, using a fully synthetic peptide of FtsB 22-87 (without FITC) produced crystals that, although in a different space group, show exactly the same structure and arrangement as observed before when adding the entire periplasmic domain of FtsB (22-103) to FtsQ periplasmic domain. F) Superposition of un-bound *E. coli* FtsQ periplasmic domain (PDB ID 2VH1) (19) and our FtsQ structure in complex with FtsB. Overall, there are only small deviations, but Y248 dramatically changes its sidechain conformer (and the entire loop 247-252 changes conformation slightly), as does W256.

Resolving only residues 64-87 in the crystal structure of the FtsQ:FtsB complex raised the question of what the remaining residues present during crystallisation do. For this we went back to SPR and tested two FtsB subdomains comprising residues 22-64 and 64-87, chemically synthesised as peptides (Suppl. Figure S2, replacing methionine 77 with norleucine). SPR analysis unequivocally showed that the region within FtsB N-terminal to the structure, residues 22-64, does not bind to FtsB, whereas the region containing only the ordered parts of the structure, residues 64-88, produced binding curves and binding parameters very similar to the original periplasmic 22-103 construct (Figure 3B & C). The binding of the 64-87 peptide was further investigated by fluorescence polarisation, a solution assay. For this, FITC-labelled FtsB 64-87 peptide was synthesised (Suppl. Figure S2) and the decrease in polarisation measured while FtsQ was added (Figure 3D). The resulting K_d_ of ~ 9 µM is less tight than what we measured by SPR and since identical reagents were used, gives an impression of the variations caused by using different assay technologies, but confirms in principle that FtsB 64-87 binds FtsQ in solution. To further show this we crystallised an acetylated version of this peptide with FtsQ, resulting in essentially the same structure as observed for FtsB 22-103, despite being in a different crystallographic space group (Figure 3E, Table 1). It is noteworthy that the synthesised FtsB 64-87 peptide is highly soluble in water, at least to 10 mM.

As already mentioned, FtsB binding to FtsQ involves only few and minor changes to the conformation of FtsQ when compared to a previous unbound structure of FtsQ (PDB ID 2VH1, Figure 3F) (19). Two exceptions are that FtsQ Y248 and W256 change their side chain conformations significantly upon binding, which fits well with our data showing that FtsQ Y248 is absolutely critical for the interaction with FtsB. The tyrosine side chain of Y248 is protruding from the main structure whereas upon interaction with FtsB it shifts deeply inward providing aromatic stacking with FtsB F84 and potential hydrogen bonding to FtsB R72.

We conclude that binding of the periplasmic domain of FtsB to FtsQ, in absence of FtsL, only involves FtsB residues 64-87 as shown by the structure and this interaction can be faithfully reconstituted by using synthesised and water-soluble peptides comprising FtsB residues 64-87.

### The FtsQ:FtsB interaction in the context of bacterial cell division

Based on the FtsQ:FtsB structure (Figure 2), we investigated *E. coli* cells harbouring FtsQ and FtsB mutant proteins in order to validate the structure and to understand the contributions of various parts of the interface to the ability to progress cell division. Functioning of the mutants was tested by low-level un-induced expression in *E. coli* strain LMC531, an *ftsQ* temperature-sensitive mutant. Cells grown at the permissive temperature were imaged by phase contrast microscopy (Figure 4A) or spotted on solid medium (Suppl. Figure S3A), followed by incubation at the permissive and non-permissive temperatures. Under these conditions, phase contrast microscopy of LMC531 cells harbouring the empty vector (EV) showed filamentation upon growth in liquid LB medium at the non-permissive temperature whereas positive control cells harbouring a tagged but otherwise wild-type FtsQ construct showed normal cells also at the non-permissive temperature (Figure 4A). Equally, in the spot assay, the negative control cells harbouring the empty vector only grew to high dilutions at the permissive temperature, whereas the positive control showed good growth at high dilution at the non-permissive temperature (Suppl. Figure S3A).

**Figure 4.**
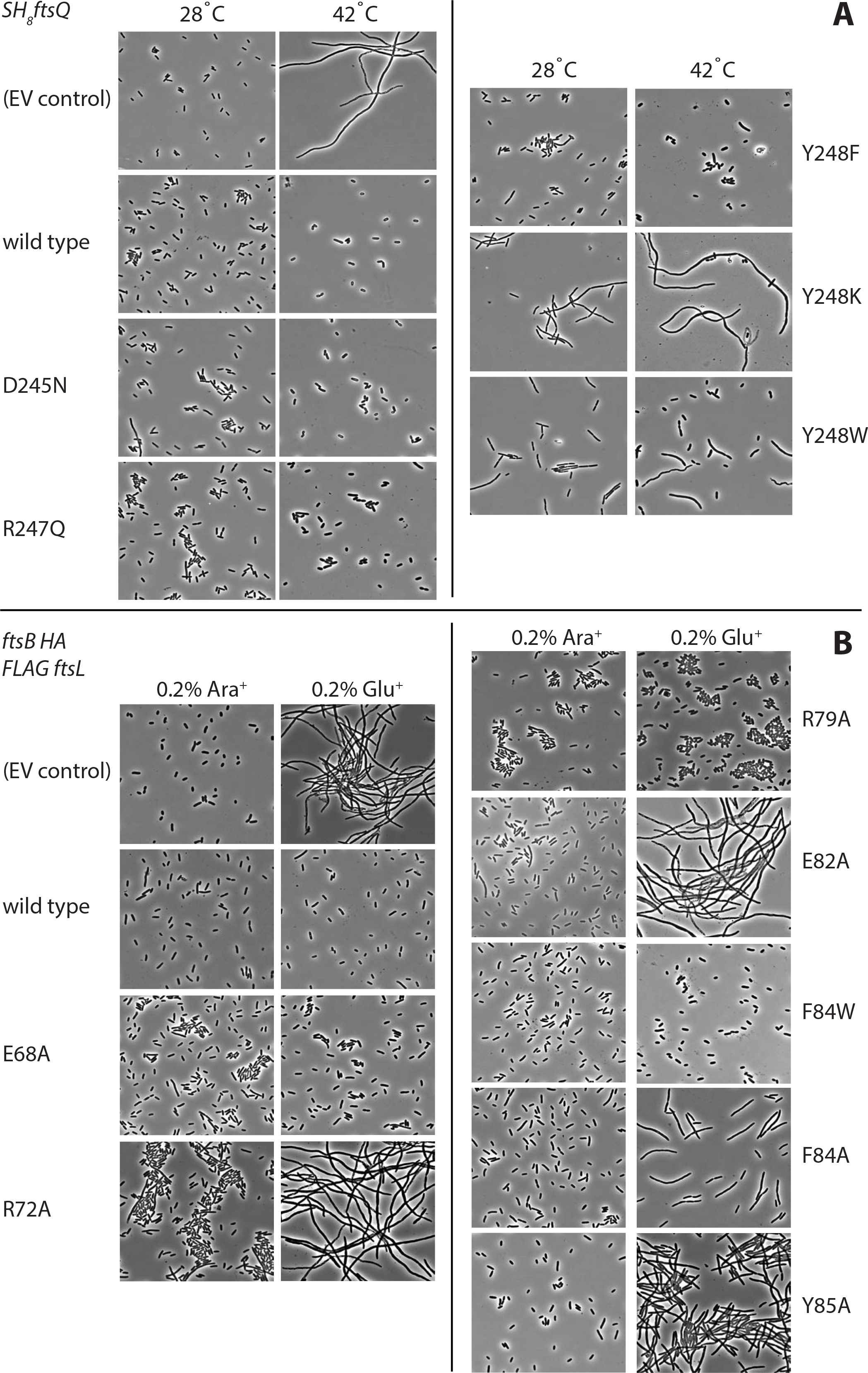
Mutating residues implicated in FtsB and FtsQ complex formation impairs cell division. A) FtsQ mutants. The FtsQ temperature-sensitive *E. coli* strain LMC531 (40), harbouring a plasmid for the expression of mutant SH_8_FtsQ was grown for 5 h at the permissive temperature (28 °C) or at non-permissive temperature (42 °C). Cells were analysed by phase-contrast microscopy. EV: empty vector/plasmid. B) FtsB mutants. The *E. coli* FtsB depletion strain NB946 (41), harbouring a plasmid for the expression of mutant FtsB-HA derivatives was grown under non-depleting conditions in the presence of 0.2% L-arabinose (Ara^+^) and under depleting conditions in the presence of 0.2% L-glucose (Glu^+^). Cells were analysed by phase-contrast microscopy. EV: empty vector/plasmid. Mutated residues are highlighted in Figure 2D.

Using both assays, mutant FtsQ Y248F showed full complementation, indicating that the hydroxyl moiety in the tyrosine side chain is not required for functioning. In contrast, FtsQ Y248K and even a Y248W substitution were unable to grow on plates at the non-permissive temperature and showed a strongly filamentous phenotype using microscopy at the non-permissive, and even at the permissive temperature, indicating a dominant negative effect on cell division. We also investigated FtsQ D245N and R247Q, conservative mutations probing the role of the polar interactions FtsB is making with FtsQ towards its N-terminal part of the binding region, and these showed no effects. Equally, FtsQ mutants S250A, G251A and W256A showed no obvious effects in the spot assay, ruling out their significance for the FtsB interaction (Suppl. Figure S3A). In order to demonstrate that phenotypes observed were not due to reduced protein levels of the mutated proteins, we performed Western blotting (Suppl. Figure S3B). Given the central role of Y248 and its very strong and selective phenotypes depending on the replacing residue types we also investigated the localisation of a fluorescently labelled version of FtsQ Y248W. This mutant protein showed strong cell division inhibition in both assays but localised normally in cells, being recruited correctly to division sites (Suppl. Figure S4), indicating that only downstream divisome interactions were affected, as predicted.

The central role of FtsQ Y248 in the interaction interface suggests that specific mutations at this position are not tolerated because they affect the interaction with FtsB. If this is the case, complementary mutations in this area of the FtsQB complex in FtsB should have a similar effect. To examine this, structure-guided FtsB mutant proteins were tested for functionality by low-level background expression in a conditional *E. coli* FtsB mutant strain NB946 in which the chromosomal *ftsB* gene is under control of an arabinose promoter. Depletion of FtsB occurs when 0.2% L-arabinose is replaced by 0.2% L-glucose in the growth medium leading to filamentation, and eventually cell death, due to the essential nature of FtsB as observed by phase contrast microscopy and a spot assay (empty vector ‘EV’ negative control, Figure 4B; Suppl. Figure S5). It also confirmed that an HA-tagged FtsB construct that was used as basis for the mutagenesis was able to complement growth and proper cell division in cells grown in the absence of the inducer L-arabinose in contrast to NB946 cells harbouring the empty vector (Figure 4B; Suppl. Figure S5). Mutations were introduced changing conserved residues within the FtsB domain binding to FtsQ and analysed using the microscopy and spot assay described above for FtsQ mutants. FtsB F84A, also compromised in the SPR assay (Figure 3A & C), appeared non-functional, although cell filamentation was relatively mild, suggesting that the aromatic interaction with FtsQ Y248 is critical for FtsB functioning. In agreement with this supposition, changing F84 into tryptophan did not affect FtsB functioning. Y85A was also non-functional with a rather strong filamentation phenotype that was even more pronounced and dominant in a (F84A, Y85A) double mutant (spot assay only, Suppl. Figure S5).

The loop between the α-helix and β-strand of the FtsB domain that is resolved in the FtsQB complex structure is connected via two salt bridges, R72-E82 and E68-R79 (Figure 2D). In agreement with the SPR data (Figure 3A & C), both R72A and E82A did not complement FtsB depletion and showed a strong filamentation phenotype suggesting that this salt bridge is essential for interaction or possibly also shaping FtsB into the correct fold for binding. In contrast, FtsB E68A and R79A appeared fully functional, suggesting this second salt bridge is largely dispensable. These findings are further supported by Suppl. Figure S1, which shows that the R72-E82 salt bridge is more conserved than E68-R79. Similarly, FtsB E65 and E69, potentially interacting with FtsQ R196, could be changed into alanine without functional consequences (spot assay only, Suppl. Figure S5). Because of the very low amount of FtsB in cells and the low protein amounts needed to complement in our assays, we have been unable to test protein levels in cells (Figure 4B and Suppl. Figure S5) by Western blotting reliably.

## Discussion

Our investigation of the interactions within the FtsQLB complex produced two surprises. It has been proposed that FtsL and FtsB form a coiled coil/leucine zipper heterodimer that binds to FtsQ (22, 23). We show here that at least the periplasmic domains of *E. coli* FtsL and *E. coli* FtsB do not interact, but both proteins bind well to FtsQ, forming a heterotrimeric complex. This has been reported before based on pull-down experiments (25). Based on these data and the lack of coiled coil/leucine zipper prediction for the periplasmic domain of FtsL, we suggest that FtsL binds to both FtsB and FtsQ in the complex, at least in *E. coli*. It is worth noting that FtsL proteins from other organisms do show weak coiled coil/leucine zipper predictions and FtsL from *Bacillus subtilis* shows strong prediction. It is conceivable that different interaction modes exist between these small proteins.

The second surprise was that the structure of the complex between the periplasmic domains of *E. coli* FtsB and FtsQ contained FtsB residues 64-87 only. We showed that this subdomain is indeed necessary for the interaction but is also sufficient, suggesting that the remaining residues within FtsB are disordered in the crystals and do not bind to FtsQ.

Overall, the interaction interface between FtsQ and FtsB observed here in the structure (Figure 2) corresponds to the main interaction site that was identified by *in vivo* site-specific photo crosslinking (24). In that study, 50 surface-exposed positions in the periplasmic domain of *E. coli* FtsQ were changed into Bpa using amber suppressor technology for photon-induced crosslinking of FtsQ with its binding partners. Using this scanning approach, the strongest crosslinking to FtsB was observed at Y248Bpa and S250Bpa. Moreover, paired cysteine mutagenesis enabled the formation of a disulfide bond between FtsQ S250C and FtsB V88C. Both findings are in line with our structure. Although FtsB 88V is not included in the structure, the last resolved residue, L87, is immediately juxtaposed to FtsQ S250 and Y248 in FtsQ is the central residue of the hydrophobic patch on FtsQ that binds FtsB. Similarly, functional analysis of the FtsQ Bpa mutants showed that they all complemented the function of FtsQ in an FtsQ Ts mutant grown at the non-permissive temperature, except for Y248Bpa, highlighting again the crucial nature of this position that does not allow even small alterations. It is also notable that the orientation of the Y248 side chain is different in the unbound FtsQ (PDB ID 2VH1) (19) as compared with the FtsQ:FtsB complex described here (Figure 3F).

Of note, FtsQ S250 and Y248 are part of a conserved region in the interaction interface with FtsB that stretches from Y243 to W256 (compare Figure 2C & D with Figure 5B). Single substitutions in this motif, D245N, R247Q, G251A, W256A (this study), G255C and S250C (24) did not affect functioning of FtsQ consistent with the observed tolerance of FtsQ towards Bpa substitutions. In an independent study, FtsQ amino acids 257 to 276 were shown to be dispensable for function whereas shorter truncations were less stable, possibly because FtsB and/or FtsL were not recruited (28).

**Figure 5.**
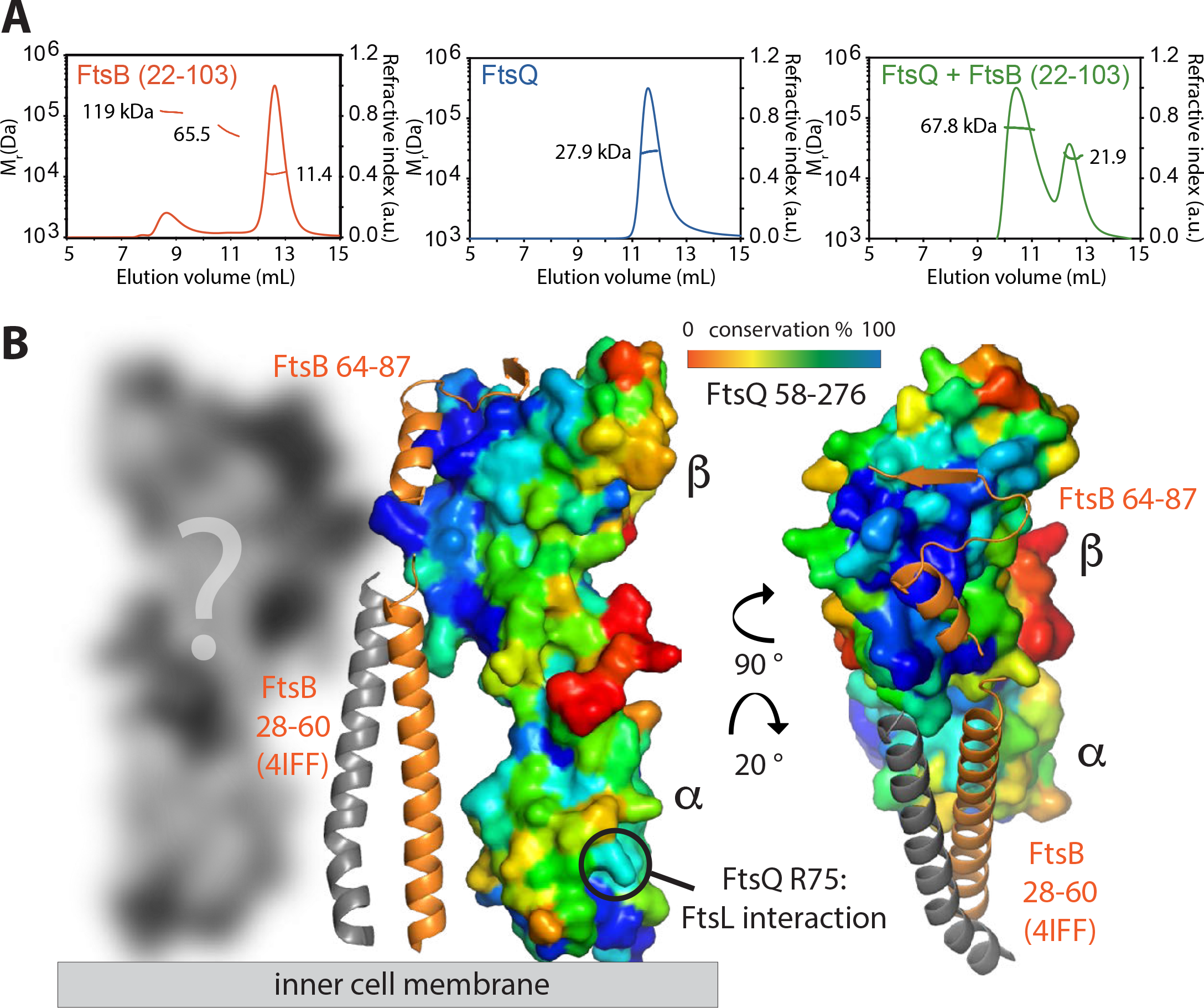
Model of the periplasmic complex between *E. coli* FtsB and FtsQ. A) Size exclusion chromatography with multiple angle light scattering (SECMALS) of the complex of FtsB and FtsQ. FtsB forms large oligomers on its own. FtsQ is monomeric alone but together, FtsB and FtsQ most likely form a 2 + 2 complex. Analytical ultracentrifugation (AUC) was also used to investigate the same complex with very similar results (Suppl. Figure S6). B) Residues 22-64 of FtsB were shown to not interact with FtsQ (Figure 3B & C). A previous crystal structure (PDB ID 4IFF) (30) showed residues 28-60 to be able to form a coiled coil arrangement with each other and we think it is likely that this interaction leads to the dimerisation of the FtsB and FtsQ complex into the observed 2 + 2 stoichiometry. Residues 22-64 within FtsB link its single transmembrane helix to the FtsQ-interacting domain in FtsB and also produce the putative dimer as shown by forming a homodimeric coiled coil. It was previously shown that a region around R75 in FtsQ links to FtsL (24). FtsQ is shown in surface representation with sequence conservation colour coded (blue: most conserved, red: least conserved). It is clear from the plot that the FtsB binding region (residues 64-87) covers most of the highly conserved patch on the β domain of FtsQ.

In the FtsQ:FtsB structure, FtsB Y85 is oriented towards FtsQ L226, a position that was shown to crosslink very strongly to FtsB previously (24). In accordance with these data, it has been shown that FtsB truncated at Y85 is unable to complement and interact with FtsQ whereas FtsB truncated at D90 is fully functional (29), and this is also supported by our data showing that FtsB 64-88 binds as well as the entire periplasmic domain comprising residues 22-103 (Figure 3B & C).

The three residues upstream of FtsB R79 have been changed individually into cysteine (S76C, M77C and T78C) in a previous study (24) without any effect on FtsB functioning, arguing that the precise sequence of the loop that connects the FtsB α-helix and β-strand is not very relevant for FtsB functioning.

Finally, we would like to speculate on the nature of the FtsQ:B complex when in the membrane and with FtsL and N. Here, we found that FtsQ and FtsB together form molecular species bigger than a 1:1 complex (Figure 2A & B). In order to understand this better, we also performed size exclusion chromatography with multiple angle light scattering (SEC-MALS) and analytical ultracentrifugation (AUC, Figure 5A, Suppl. Figure S6A). Size estimates from those experiments for FtsB, FtsQ and FtsQ:B are summarised in Suppl. Figure S6B. FtsB alone occurred in several species, including some high molecular weight oligomers. FtsQ was almost exclusively monomeric and the FtsQ:FtsB complex showed two species, most likely corresponding to 1:1 and 2:2 complexes. Taken this into account and the previous structure of the coiled coil segment of FtsB comprising residues 28-60 (PDB ID 4IFF) (30), a hybrid model of the entire periplasmic FtsQ:FtsB complex can be assembled (Figure 5B). In this model, the FtsB coiled coil segment, disordered in our crystal structure, forms a homodimer, as observed in the 4IFF crystal structure (that, admittedly, was artificially dimerised by fusing it to a coiled coil dimerisation domain) (30). FtsQ only binds to the C-terminal part of FtsB, containing residues 64-87 as demonstrated here by the structure and other experiments. Because FtsB is a dimer, this means that the complex will recruit two FtsQ molecules, explaining the observed 2:2 stoichiometry and a measured molecular mass of 68-69 kDa. The model also predicts that the transmembrane segments of FtsB are dimerised or in very close proximity, in contrast to those of the two FtsQ molecules, that could be further apart. As shown, FtsL binds together with FtsB to FtsQ, most likely binding to surfaces on FtsB and FtsQ simultaneously. It is not clear from looking at sequence conservation on the FtsQ surface where those binding surfaces are located since the FtsB binding covers almost perfectly a highly conserved patch on FtsQ (Figure 5B, blue patch), although we would like to speculate that FtsL might bind to the second interaction hotspot on FtsQ around residue FtsQ R75 (24). Equally, FtsN binding, demonstrated here biochemically, involves binding to FtsQ independently of FtsB and the location of that binding site is currently unknown.

With our crystal structure we conclude that it is a valid representation of the complex formed during cell division and relates to previous data regarding FtsQ and FtsB. The complex represents an exciting target for structure-based design of protein interaction inhibitors shutting down cell division based on peptides or peptide mimetics as the FtsQ:FtsB interaction surface is small, the 64-87 FtsB peptide is water soluble and the interaction is easily interrupted by single mutations. In future, it may be possible to biochemically add more components of the divisome as we report here direct interactions of FtsQ also with FtsL and FtsN, and to investigate the structure and function of divisome subcomplexes in the current absence of any reports of isolated or over-expressed divisome holo complexes, with only one study showing high-molecular weight divisome complexes by Western blotting (31).

## Materials and Methods

### Surface plasmon resonance (SPR)

SPR was performed using a Biacore T200 instrument using CM5-sensor chips (GE Healthcare). Both reference control and analyte channels were equilibrated in HBS-N buffer (0.01 M Hepes pH 7.4, 0.15 M NaCl) at 20 °C. FtsQ (58-276) or FtsL (58-121) were immobilised onto chip surfaces through amide coupling using the supplied kit to reach RU values (resonance units) of ~1000 and 3300, respectively.

Triplicate SPR runs were performed in 10 mM Hepes, pH 7.4, 150 mM NaCl, 0.4 % (w/v) glycerol with analytes injected for 120 s followed by a 600 s dissociation in 1:2 dilution series with initial concentrations of FtsB (22-103), FtsB (64-88) of 80 µM; FtsB (22-64) of 200 µM; FtsB-F84A and FtsB-R72A of 68 µM; FtsN (58-320) and FtsN (58-320) + FtsB (22-103) of 40 µM each for experiments where FtsQ was the ligand. For runs where FtsL (58-121) was attached to the chip surface, initial concentrations of FtsQ (58-276) and FtsB (22-103) were 40 µM each either alone or in combination. The surface was regenerated with 10 mM HCl for 60 s. After reference and buffer signal correction, sensogram data were fitted using KaleidaGraph (Synergy Software) and Prism (GraphPad Software).

### Cloning, protein expression and purification

FtsQ (58-276) and FtsN (57-319) from *Escherichia coli* strain K12 (FtsQ: FTSQ_ECOLI; FtsN: FTSN_ECOLI) were amplified using PCR from genomic DNA, cloned into the NdeI/BamHI sites of plasmid pHis17 and expressed as a C-terminal His_6_-tag in *E. coli* (see Table 1 for exact protein sequences).

FtsB (22-103) and FtsL (58-121) from *E. coli* (FtsB: FTSB_ECOLI; FtsL: FTSL_ECOLI) were amplified using PCR from genomic DNA, cloned into the NdeI/BamHI sites of plasmid pET15b and expressed as a N-terminal His_6_-tag in *E. coli* (see Table 1 for exact protein sequences).

Recombinant proteins were expressed in *E. coli* C41(DE3) cells, which were grown in 2xTY media with ampicillin (100 mg/L) at 37 °C with 200 rpm shaking to an OD_600_ of 0.6. Cultures were then shifted to 20 °C and induced by the addition of 0.5 mM isopropyl-β-D-1-thiogalactopyranoside (*IPTG*), before overnight incubation. Selenomethionine-substituted FtsB (22-103) cell cultures were grown to early log phase (A_600_ = 0.6) at 37 °C in M9 minimal medium, supplemented with 0.4 % glucose, 2 mM MgSO_4_. 100 mg/L of DL-selenomethionine (Generon), 100 mg/L of lysine, threonine, phenylalanine and 50 mg/l of leucine, isoleucine and valine were added as solids. Fifteen minutes later, protein expression was induced with 0.5 mM IPTG and grown overnight at 20 °C.

### Protein purification

For crystallisation experiments, bacterial cell pellets containing over-expressed FtsQ (58-276) and FtsB (22-103) were mixed and resuspended in 50 mM Tris, 150 mM NaCl, 10 % glycerol, pH 8.0. Cell lysis was carried out at 25 kPSI using a cell disruptor (Constant Systems) and the lysate was clarified by centrifugation at 20,000 rpm for 30 min at 4 °C. The supernatant was passed over a HisTrap™ HP column (GE Healthcare). The column was equilibrated with 25 mM Tris, 150 mM NaCl, 10 % glycerol, 0.5 mM DTT, pH 8.0. Proteins were eluted with 300 mM imidazole in the same buffer. Peak fractions were concentrated and loaded onto a HiLoad Sephacryl S300 16/60 column (GE HEalthcare) equilibrated in 25 mM Tris, 100 mM NaCl, 2 mM DTT, pH 7.4. Purified FtsQB complex was concentrated to 6 mg/mL using centrifugal concentrators (Vivaspin, Sartorius) for immediate use. For SPR and SEC-MALS assays, FtsB, FtsL, FtsQ and FtsN proteins were purified separately using the same protocol, then concentrated to 5-10 mg/mL before freezing in liquid nitrogen and storage at −80°C.

### *Crystallisation*, data collection and structure determination

Crystallisation conditions were found using our in house high-throughput crystallisation platform (32), by mixing 200 nL *SeMet*-substituted FtsBQ or wild type FtsQB solution at 6 mg/mL, with 200 nL of 1920 different crystallisation reagents in MRC vapour diffusion sitting drop plates. Crystals were grown at 19 °C by vapour diffusion in 0.15 M potassium thiocyanate, 20 % PEG 550 MME, 0.1 M sodium cacodylate pH 6.5.

For the FtsQ (58-276)/FtsB (64-88 peptide) complex crystallisation, FtsB synthetic peptide was added to 4 mg/mL FtsQ protein solution with molar ratio of 1 FtsQ:4 FtsB. Crystals were grown at 19 °C by vapour diffusion in 0.057 M potassium thiocyanate, 10 % PEG 550 MME, 0.1 M sodium cacodylate pH 6.5. 25 % (v/v) glycerol was used as a cryo-protectant. Conditions yielding crystals were optimised, and crystals from either the initial screens or subsequent optimisations were selected for data collection. Diffraction images were collected from single frozen crystals at beamlines I03, I04 and I04-1 at Diamond Light Source (Harwell, UK) as indicated in Table 1. Diffraction images were processed with XDS (33) and further processed with the CCP4 package of programs (34). Initial phases were determined by molecular replacement using PHASER with a previous *E. coli* FtsQ structure as search model (PDB ID 2VH1) (35). Iterative model building and refinements were carried with MAIN, REFMAC and PHENIX (36–38). Data and model statistics are summarised in Table 1.

### FtsB (64-87) peptide synthesis, purification and characterisation

Peptides were synthesised by Fmoc-based solid-phase peptide synthesis on H-rink amide ChemMatrix^®^ resin (Merck). The peptide sequence was assembled using an automated synthesiser (Syro II, MultiSynTech). For amino acid coupling 4 equivalents (eq) of the Fmoc protected amino acids (Iris Biotech) according to the initial loading of the resin were mixed with 4 eq of 1-[bis(dimethylamino)-methylene]-1*H*-1,2,3-triazolo[4,5-*b*]pyridinium 3-oxid hexafluorophosphate (HATU) and 8 eq *N*,*N*-diisopropylethylamine (DIPEA) and added to the resin for 40 min. In a second coupling step, the resin was treated with 4 eq of the Fmoc-protected amino acid mixed with 4 eq benzotriazole-1-yl-oxy-tris-pyrrolidino-phosphonium hexafluorophosphate (PyBOP) and 8 eq 4-methylmorpholine (NMM) for 40 min. After double coupling a capping step to block free amines was performed using acetanhydride and DIPEA in *N*-methyl-2-pyrrolidinone (NMP) (1:1:10) for 10 min. Fmoc deprotection was performed using 20 % piperidine in dimethylformamide (DMF) for 5 minutes, twice. After each step the resin was washed 4 times with DMF. For the crystallisation experiment of FtsB 64-87 with FtsQ, the peptide was acetylated with Ac_2_O final Fmoc deprotection. The fluorescently labelled peptide for affinity measurements was *N*-terminally coupled with 8-(9-fluorenylmethyloxycarbonyl-amino)-3,6-dioxaoetanoic acid as stated above, and subsequently fluoresceinisothiocyanat (FITC) was coupled manually using 8 eq DIPEA for 1.5 h, twice. Final cleavage was performed with 94 % trifluoroacetic acid (TFA), 2.5 % 1,2-ethanedithiole (EDT), 2.5 % H_2_O and 1 % triisopropylsilane (TIPS) for 1.5 h, twice. The cleavage solutions were combined and peptides were precipitated with diethyl ether (Et_2_O) at −20 °C for 10 min. Peptides were resolved in water/acetonitrile (ACN) 5:5 and purified by reversed-phase high performance liquid chromatography (HPLC; Nucleodur C18 column [Macherey-Nagel]; 10×125 mm, 110 Å, 5 µm particle size) using a flow rate of 6 mL/min (A: ACN with 0.1 % TFA, B: water with 0.1 % TFA). Obtained pure fractions were pooled and lyophilised. Peptide characterisation was performed by analytical HPLC (1260 Infinity [Agilent Technology]; flow rate of 1 mL/min, A: ACN with 0.1 % TFA, B: water with 0.1 % TFA) coupled with a mass spectrometer (6120 Quadrupole LC/MS [Agilent Technology]) using electro spray ionisation (Eclipse XDB-C18 column [Agilent], 4.6×150 mm, 5 µm particle size). Analytical HPLC chromatograms recorded at 210 nm are shown in Suppl. Figure S2. Quantification of acetylated peptide was performed by HPLC-based comparison (chromatogram at 210 nm) with a reference peptide and quantification of fluorescein labelled peptide was performed using the extinction coefficient ε=77.000 M^−1^cm^−1^ of fluoresceinisothiocyanat (FITC) in 100 mM sodium dihydrogen phosphate; pH 8.5.

### Fluorescence Polarisation Assay (FP)

To determine the affinity of the FtsB peptide to FtsQ, a 0.1 mM DMSO solution of the FITC-labelled peptide was dissolved in 10 mM Hepes pH 7.4, 150 mM NaCl, 0.1% Tween-20 to yield a 40 nM peptide solution. A 3-fold dilution of FtsQ (15 μL per well) was presented in a 384-wellplate (black, flat bottom [Corning]) and incubated with the peptide solution (5 μL, final peptide concentration 10 nM). The final protein concentration ranged from 800 µM − 0.5 nM. After incubation for 1 h at room temperature, fluorescence polarisation was measured using a Tecan Spark 20M plate reader with λ_ex_ = 485 nm and λ_ex_ = 525 nm. K_d_-values were determined by nonlinear regression analysis of dose-response curves using GraphPad Prism software.

### Size exclusion chromatography with multi-angle light scattering (SEC-MALS)

The masses in solution of FtsB (22-103), FtsQ (58-276) both single and in a complex were estimated using an online Dawn Heleos II 18 angle light scattering instrument (Wyatt Technologies) coupled to an Optilab rEX online refractive index detector (Wyatt Technologies). Protein samples (100 µL) were resolved on a Superdex S-75 10/300 analytical gel filtration column (GE Healthcare), pre-equilibrated with 25 mM Tris pH 7.4, 100 mM NaCl, 1 mM DTT at 0.5 mL/min. Protein concentration was determined from the excess differential refractive index based on dn/dc of 0.186 mg/mL. The light scattering and protein concentration at each point across the peaks in the chromatograph were used to determine the absolute molecular mass from the intercept of the Debye plot using Zimm’s model as implemented in the ASTRA version 5.3.4.20 software (Wyatt Technologies). In order to determine the inter-detector delay volumes, band broadening constants and the detector intensity normalisation constants for the instrument, we used BSA as a standard prior to sample measurements.

### Analytical Ultracentrifugation

Samples of FtsB (22-103), FtsQ (58-276) alone or mixed together at concentrations of 1 mg/mL were subjected to velocity sedimentation at 50,000 rpm at 20 °C in 25 mM Tris pH 7.4, 100 mM NaCl, 1 mM DTT using 12 mm double sector cells in an An50Ti rotor using an Optima XL-I analytical ultracentrifuge (Beckmann). The sedimentation coefficient distribution function, c(s), was analysed using the SEDFIT program, version 15.0 (39). The partial-specific volumes (v-bar), solvent density and viscosity were calculated using SEDNTERP software (SEDNTERP: Thomas Laue, University of New Hampshire).

### Plasmids, strains and growth conditions for complementation assays

*E. coli* strains BL21(DE3), LMC531 (*ftsQ1*[Ts]) (40), and NB946 (41) were grown in LB medium with shaking at 200 rpm. When indicated, strains LMC531 and NB946 were grown in GB1 medium (4.83 g of K_2_HPO_4_· 3H_2_O, 2.95 g of KH_2_PO_4,_ 1.05 g of (NH_4_)_2_SO_4_, 0.10 g of MgSO_4_ · 7H_2_O, 0.28 mg of FeSO_4_ · 7H_2_O, 7.1 mg of Ca(NO_3_)_2_ · 4H_2_O, 4 mg of thiamine, 0.4% glycerol [v/v], 0.2% casamino acids [w/v] per litre at pH 7.0). When required, 0.2% L-arabinose, 0.2% L-glucose, 100 µg/mL ampicillin, 30 µg/mL chloramphenicol, 25 µg/mL kanamycin, and 50 µg/mL spectinomycin were added to the culture medium.

Standard PCR and cloning techniques were used for DNA manipulation. p29SENX-SH8FtsQ encoding full length FtsQ with an amino-terminal StrepII-His8 dual tag (24) was used as template for mutagenesis in *SH8ftsQ* using overlap extension PCR.

The DNA sequence encoding full length FtsB with a carboxy-terminal HA tag and the DNA sequence encoding full length FtsL with an amino-terminal FLAG tag were cloned by PCR and transferred into the pCDF plasmid (Novagen) resulting in pCDF-FtsB-HA FLAG-FtsL. Mutations in *ftsB-HA* were introduced by overlap extension PCR.

Plasmids pTHV037-mNG-FtsQ was constructed by cloning *ftsQ* into pTHV037 (42). *ftsQ* was amplified from *E. coli* genomic DNA with primers containing restriction sites for *Eco*RI and *Nco*I, also used to digest pTHV037. A triple asparagine linker was introduced immediately downstream the *Eco*RI site. Plasmid and insert were ligated with T4 DNA ligase (NEB, Ipswich, MA). Finally, the plasmid was amplified by circular PCR to incorporate the Y248W mutation and obtain the plasmid pTHV037-mNG-FtsQY248W.

### Functionality and detection of FtsQ and FtsB mutants

To assess functioning of FtsQ mutants by microscopy, *E. coli* LMC531 cells harbouring one of the p29SENX-SH8FtsQ mutants were grown overnight at 28°C in LB medium supplemented with 100 µg/mL ampicillin. The same medium was inoculated 1:100 with the overnight cultures and incubated for 5 h at 28°C or 42°C. To assess functioning of FtsB mutants by microscopy, *E. coli* NB946 cells harbouring one of the pCDF-FtsB-HA FLAG-FtsL mutants were grown overnight at 37 °C in LB medium supplemented with 25 µg/mL ampicillin, 20 µg/mL kanamycin and 0.2% L-arabinose. Then, LB medium supplemented with the indicated antibiotics and 0.2% L-glucose or 0.2% L-arabinose was inoculated 1:100 with the overnight cultures and incubated for 5 h at 37 °C. The cells were fixed by addition of formaldehyde to 3% at room temperature for 15 min, harvested by centrifugation (13,000 x *g*, 2 min) and resuspended in PBS for the examination of cell morphology by phase-contrast microscopy. To determine the expression of mutant FtsQ under these conditions, cells were harvested from the cultures by centrifugation (13,000 x *g*, 2 min) and the pellet was resuspended in 2X SDS sample buffer and incubated for 5 min at 96 °C. The crude cell lysates of LMC531 were separated on 12% SDS-PAGE gels and analysed by Western blotting using anti-FtsQ and anti-FtsB affinity-purified rabbit polyclonal antibodies.

To assess functioning of FtsQ mutants by complementation of growth on solid medium, *E. coli* LMC531 cells harbouring one of the p29SENX-SH8FtsQ mutants were grown overnight at 28 °C in LB medium with 100 µg/mL ampicillin. The same medium was inoculated 1:100 with the overnight cultures, incubated for 10 h at 28°C and then diluted 1:100 in GB1 medium with 100 µg/mL ampicillin for overnight growth at 28°C. The same medium was inoculated 1:100 with the overnight cultures and incubated for 5 h at 28 °C or 42 °C. The cultures were ten-fold serially diluted and 4 µl of each dilution was spotted on GB1 agar medium. The growth was assessed after 21 h incubation at 28°C or 18 h at 42 °C.

To assess functioning of FtsB mutants by complementation of growth on solid medium, *E. coli* NB946 cells harbouring one of the pCDF-FtsB-HA mutants were grown overnight at 37 °C in LB medium supplemented with 25 µg/mL ampicillin, 20 µg/mL kanamycin and 0.2% L-arabinose. The same medium was inoculated 1:100 with the overnight cultures and incubated for 10 h at 37 °C and then diluted 1:100 in the same GB1 medium for overnight growth at 37 °C. The same medium was inoculated 1:100 with the overnight cultures and incubated for 5 h at 37 °C. The bacterial suspensions were diluted and spotted as described above on GB1 agar medium supplemented with the indicated antibiotics and either 0.2% L-arabinose or 0.2% L-glucose and growth was analysed after 18 h incubation at 37°C.

### Fluorescence microscopy

Temperature sensitive FtsQ strain LMC531 (40) containing plasmid pTHV037-mNG-FtsQ or pTHV037-mNG-FtsQY248 was grown in LB overnight at 30 °C supplemented with 100 µg/mL ampicillin. The day after, samples were diluted (1:500) in 25 mL of the same medium and grown at 30 °C until OD_600_ was 0.25. Cells were then diluted 1:10 in cultures at 30 or 42 °C. When OD_600_ was 0.05, samples were induced with 15 µM IPTG for 2 mass doubling times. Non-induced samples were used as control. Cells were immobilised on 1 % agarose (43) and imaged with a Nikon Eclipse T1 microscope (Nikon Plan Fluor 100x/1.30 oil ph3 DLL objective) coupled to an EMCCD camera (Hamamatsu Flash 4.0).

## Acknowledgements

We would like to thank Gregory Koningstein (VU University) for technical help, Minmin Yu (MRC-LMB) for help with crystallographic data collection and Fusinita van den Ent (MRC-LMB) for advice and discussions. AMS and TdB received support from the NAPCLI project within the JPIAMR program (ZonMW project 60-60900-98-207). This work was funded by the Medical Research Council (U105184326 to JL) and the Wellcome Trust (202754/Z/16/Z to JL).

## Figure legends Supplementary Figures

**Supplementary Figure S1.** Sequence conservation-based colouring of FtsB surface, shown in stereo. Around 100 sequences similar to *E. coli* FtsB (database ID FTSB_ECOLI) were identified using BLAST and sequence conservation was determined from a CLUSTAL OMEGA multiple sequence alignment with CONSURF (http://consurf.tau.ac.il/2016/). FtsB residues 64-87 are shown as a stick model coloured by conservation from blue (most conserved residues in the alignment) to red (least conservation). FtsQ is shown as a grey molecular surface. All residues involved in major interactions with FtsQ are conserved in FtsB, in addition to internal interactions, such as R72 - E82.

**Supplementary Figure S2.** Table: peptide characteristics (molecular formula, one-letter code sequence, molecular weight, calculated and detected m/z ratios for multi charged masspec ions ([M+nH]^n+^). Abbreviations: Ac: acetyl; FITC: fluoresceinisothiocyanat fluorophore; PEG: 8-amino-3,6-dioxaoctanoyl linker; Nle: norleucine amino acid replacing methionine 77. **A)** HPLC chromatogram at 210 nm of peptide used for co-crystallisation with FtsQ (Figure 3D), gradient: 20-50 % ACN in 10 min and mass spectra of corresponding peptide. **B)** HPLC chromatogram at 210 nm of FITC PEG peptide for fluorescence polarisation (Figure 3D), gradient: 30-60 % ACN in 10 min, starting at 3 minutes and mass spectra of corresponding peptide.

**Supplementary Figure S3.** A) Spot assay of FtsQ mutants used in Figure 4A. B) Western blot showing expression levels of FtsQ mutants used in A).

**Supplementary Figure S4.** Fluorescence microscopy localisation of FtsQ (Y248W) mutant protein in *E. coli* cells. Phase contrast images, fluorescent images and fluorescence profiles per cell (from 0 to 200 AU) plotted against normalised cell length (from 0 to 100 %) of ts FtsQ LMC531 strain expressing wild type FtsQ or FtsQY248W fused to mNeonGreen (mNG) at 30 or 42°C. *n* indicates number of cells. The graphs show the average total fluorescence per cell plotted against the normalised cell length. Filaments have therefore a higher fluorescence per cell. In addition, the expression of the proteins is higher at the non-permissive temperature. Scale bar represents 2 µm.

**Supplementary Figure S5.** Spot assay showing FtsB mutants used in Figure 4B.

**Supplementary Figure S6.** A) Analytical ultracentrifugation (AUC) of FtsQ, FtsB and in complex. B) Summary table of SEC-MALS (Figure 5A) and AUC data (panel A) describing the properties of FtsQ, FtsB and their complexes.

